# The antimicrobial potential of traditional remedies of Indigenous Peoples from Canada against MRSA planktonic and biofilm bacteria in wound-infection mimetic conditions

**DOI:** 10.1101/2024.09.08.611890

**Authors:** Colin D. Rieger, Ahmed M. Soliman, Kateryna Kaplia, Nilrup Ghosh, Alexa Cervantes Lopez, Surya Arcot Venkatesan, Abraham Gildaro Guevara Flores, Matheus Antônio Filiol Belin, Florence Allen, Margaret Reynolds, Betty McKenna, Harold Lavallee, Archie Weenie, Thomas Favel, Fidji Gendron, Vincent E. Ziffle, Omar M. El-Halfawy

**Affiliations:** Department of Chemistry and Biochemistry, Faculty of Science, University of Regina, Regina, SK, S4S 0A2, Canada; Department of Microbiology and Immunology, Faculty of Pharmacy, Kafr-Elsheikh University, Kafr El-Sheikh 33516, Egypt; Department of Indigenous Knowledge and Science, Faculty of Science, First Nations University of Canada, Regina, SK, S4S 7K2, Canada; Kingston University, Faculty of Health, Science, Social Care and Education, KT1 2EE, UK; IISER Kolkata, West Bengal 741246, India; Instituto Tecnológico y de Estudios Superiores de Monterrey, 64849 Monterrey, N.L, Mexico; Engineering and Technology, Rajalakshmi Engineering College, Kanchipuram, Tamil Nadu, 602105, India; Autonomous University of Nuevo Leon (Medicine College), Department of Chemistry, 66455 San Nicolás de los Garza, N.L., Mexico; Universidade Estadual Paulista (UNESP), Campus de Botucatu. Faculdade de Medicina (FMB), 18618-970, Brazil; Elder from Peter Ballantyne Cree Nation in Saskatchewan, Treaty 6 Territory, Canada; Elder from English River First Nation in Saskatchewan, Treaty 10 Territory, Canada; Elder from Shoal River Band in Manitoba, Treaty 4 Territory, Canada; Elder from Piapot First Nation in Saskatchewan, Treaty 4 Territory, Canada; Elder from Sweetgrass First Nation in Saskatchewan, Treaty 6 Territory, Canada; Elder from Kawacatoose First Nation in Saskatchewan, Treaty 4 Territory, Canada; Department of Microbiology and Immunology, Faculty of Pharmacy, Alexandria University, Alexandria, 21521, Egypt

**Keywords:** Wound infection-mimetic conditions, MRSA, Indigenous remedies, biofilm prevention, biofilm eradication

## Abstract

Methicillin-resistant *Staphylococcus aureus* (MRSA) is the leading cause of wound infections, often progressing into serious invasive bloodstream infections. MRSA disproportionately affects Indigenous peoples in Canada with higher rates of skin and wound infections, an example of persistent gaps in health outcomes between Indigenous and non-Indigenous peoples precipitated by the legacy of colonialism. Conversely, Indigenous peoples have long used natural remedies for infections and other diseases; however, their knowledge was rarely considered for modern medicine. The stagnant antibiotic discovery pipeline and alarming rise of resistance to current antibiotics prompted us to turn to Indigenous medicine as an untapped source of antimicrobials. As such, we collected and prepared 85 extracts of medicinal plants of value to Indigenous Peoples spanning the Canadian Prairies. We explored the antimicrobial potential of these extracts against MRSA under wound infection-mimetic conditions compared to culture media typically used to study bacterial antibiotic responses and biofilms but not adequately representative of infection sites. We identified extracts with MRSA growth inhibitory [e.g., bergamot, dock, gaillardia, and dandelion extracts] and biofilm prevention and eradication [e.g., gumweed extracts] activities. Extracts, including those of chokecherry, hoary puccoon, and Northern bedstraw, were only active under wound infection-mimetic conditions, highlighting the relevance of antibiotic discovery under host-relevant conditions. Testing growth inhibitory extracts against an *S. aureus* cross-resistance platform suggested that they act through mechanisms likely distinct from known antibiotic classes. Together, through an interdisciplinary partnership leveraging Western approaches and traditional Indigenous knowledge, we identified plant extracts with promising antimicrobial potential for drug-resistant MRSA wound infections.

## Introduction

Antimicrobial resistance (AMR) is rising at an alarming rate, posing a threat to public health and modern medicine where the worldwide death toll associated with bacterial AMR was estimated at ∼5 millions in 2019 alone (1). The AMR crisis prompted the World Health Organization to issue a list of priority pathogens posing the most serious threat that require urgent action (2, 3). Included in the list is methicillin-resistant *Staphylococcus aureus* (MRSA), which continuously develops resistance to antibiotics, including last-resort ones, and remains a high-priority pathogen due to its potentially severe infections and persistent prevalence worldwide being one of the leading causes of healthcare-associated and community-acquired infections (2, 3). MRSA is the leading cause of wound infections, posing severe clinical problems (4); ∼2% of Canadians suffer from chronic wounds with direct healthcare costs of ∼$4B each year (5).

The legacy of colonialism and the associated intergenerational trauma have led to persistent troubling gaps in health outcomes between Indigenous and non-Indigenous peoples in Canada, including rates and prognosis of infectious diseases (6). Whereas Indigenous peoples in Canada have higher rates of tuberculosis, pneumococcal disease, gastrointestinal infections, and sexually-transmitted infections (including chlamydia), none of these conditions are associated with a higher risk of resistant infections among Indigenous relative to non-Indigenous people (7). For example, rates of resistant tuberculosis infections are lower among Indigenous compared to non-Indigenous people (8). In contrast, MRSA is the main drug-resistant pathogen disproportionately affecting Indigenous peoples (7, 9). Indigenous people suffer higher rates of MRSA skin infections, mainly of wounds, which often progress into serious invasive bloodstream infections (7, 10, 11).

Despite recent advances and antibiotic discovery efforts, topical and systemic antibiotics are mostly ineffective at clearing MRSA wound infection (12). Further, MRSA may colonize chronic wounds as biofilms (4, 13, 14). Biofilms are surface-associated microbial communities embedded within a self-produced extracellular polymeric matrix (4); they contribute to pathogens’ ability to tolerate antibiotics and cause persistent infections (15, 16). Currently-available antibiotics that are active against planktonic cells – non-adherent bacteria – are mostly ineffective against biofilms (16). Most antibiotics in clinical use are synthetic small molecules and analogs of Actinomycetes-derived natural products (17). Therefore, there is an urgent need to explore previously untapped sources to discover new therapeutics that can eradicate MRSA infections.

Indigenous peoples in Canada have long used natural products derived from plants and other sources as medicine. They have used over 400 different species of plants, lichens, fungi, and algae to treat a range of illnesses (18). Such knowledge, maintained and passed across generations by Elders and other Traditional Knowledge Keepers (19), has yet to be considered in Western medicine. Traditionally, Indigenous Peoples have used plants, such as Nootka rose (*Rosa nutkana* C. Presl), to cure skin infections (20). Recent studies tested select Indigenous medicinal plants, showing efficacy against *S. aureus* and other skin pathogens (20, 21). However, these studies are not comprehensive; Indigenous medicinal plants remain mostly unexplored for their antibacterial activity.

Bacterial responses to antimicrobials and biofilm formation are often studied under standard *in vitro* conditions [e.g., in Mueller Hinton broth (MHB) composed of beef extract, casein hydrolysate, and starch (22)], which do not adequately represent the conditions at the infection site. We hypothesized that screening for new antimicrobials under wound infection-mimetic conditions will uncover agents with better potency in eradicating wound infection as they may target specific antibiotic response and virulence factors that bacteria deploy during wound infections but go undetected under standard test conditions. Indeed, culture conditions and media composition dictate bacterial gene expression and dispensability profiles to meet their survival requirements (23, 24). Wound infection-mimetic *in vitro* and *ex vivo* model systems are currently available (25–27). Notably, the high organic proteinaceous load at the wound site may alter the activity of antimicrobials at the infection site (28); these and any similar alterations may not be detected in standard *in vitro* conditions but should be captured in infection-mimetic ones. Not surprisingly, bacterial antibiotic susceptibility patterns in various infection-mimetic conditions were more often predictive of the therapeutic outcome in animal experiments or patients despite discordance with the standard culture media results (29–31). The clinical outcome of antibiotic treatment often does not correlate with antibiotic susceptibility results from these standardized testing conditions (32–34). These observations highlight the need to screen for new antimicrobials under infection-like conditions.

Herein, we explored the antimicrobial potential of Traditional Indigenous remedies against MRSA under wound infection-mimetic conditions to address MRSA wound infections as an Indigenous health priority (7, 9–11), ensuring mutual benefits and reciprocity in partnerships between Indigenous and non-Indigenous researchers. We prepared extracts of various plants used as Indigenous remedies and tested the antimicrobial properties of 85 extracts against MRSA under wound infection-mimetic conditions in comparison to standard lab culture conditions. We screened these extracts for planktonic growth inhibitory effects as well as biofilm prevention and eradication against MRSA, uncovering activities under each of these conditions with mechanisms likely distinct from that of any of the known antibiotic classes. Together, through this interdisciplinary partnership that leverages Western approaches and traditional Indigenous knowledge to co-create new antimicrobial solutions, we identified medicinal plant extracts with promising antimicrobial potential for drug-resistant MRSA wound infections.

## Results and discussion

### Collection of Indigenous remedies and extract preparation

Through guidance from Elders and Traditional Knowledge Keepers with experience in Medicine Walks and picking sessions and Knowledge on Traditional medicinal plants, we collected an array of Indigenous medicinal plants from diverse ecoregions within three ecozones (prairie, boreal plain, and boreal shield). These picking sites spanned Treaties 4, 6, and 10 Territories located in the Canadian Prairies, predominantly in the Canadian Province of Saskatchewan. Each ecozone harbors specific assortments of plants due to differences in soil type and climate. Elders and Traditional Knowledge Keepers guided the identification and harvesting of different parts of the plants and the best season to harvest each plant. In total, we have collected and identified 46 plant part specimens, belonging to 28 species. Rather than exploring remedies traditionally used for infections, we will also test plants used by Indigenous Peoples for other indications akin to the current trend of repurposing existing drugs as antimicrobials. Importantly, we follow culturally appropriate protocols while respecting the First Nations principles of OCAP^®^ (35), such as recognizing the harvesting land and offering tobacco to the land and Traditional Knowledge Keepers and honouraria as appropriate. We conducted taxonomic identification of plant materials using theplantlist.org and worldfloraonline.org, in which a consortium of leading botanical institutions worldwide lists all known plant species.

Next, we prepared a total of 85 extracts from the different parts of the harvested plants separately using multiple established extraction protocols in solvents that varied from polar [water, ethanol, and methanol] to non-polar [hexane] (20); Table 1 shows the extraction details of extracts of interest. We posit that multiple preparation protocols, plant parts, and solvents will yield diverse active components possessing various biological activities. We then dried the extracts under vacuum using DNA 110 SpeedVac® concentrator to eliminate any potential antimicrobial activity of the extraction solvents and to facilitate comparison between the different extracts; the dry weight of each was later used to determine the concentrations as % w/v in the dose-response assays. We reconstituted all dry masses in a fixed volume of DMSO and stored them at −80°C, creating our stock screening library.

**Table 1.**
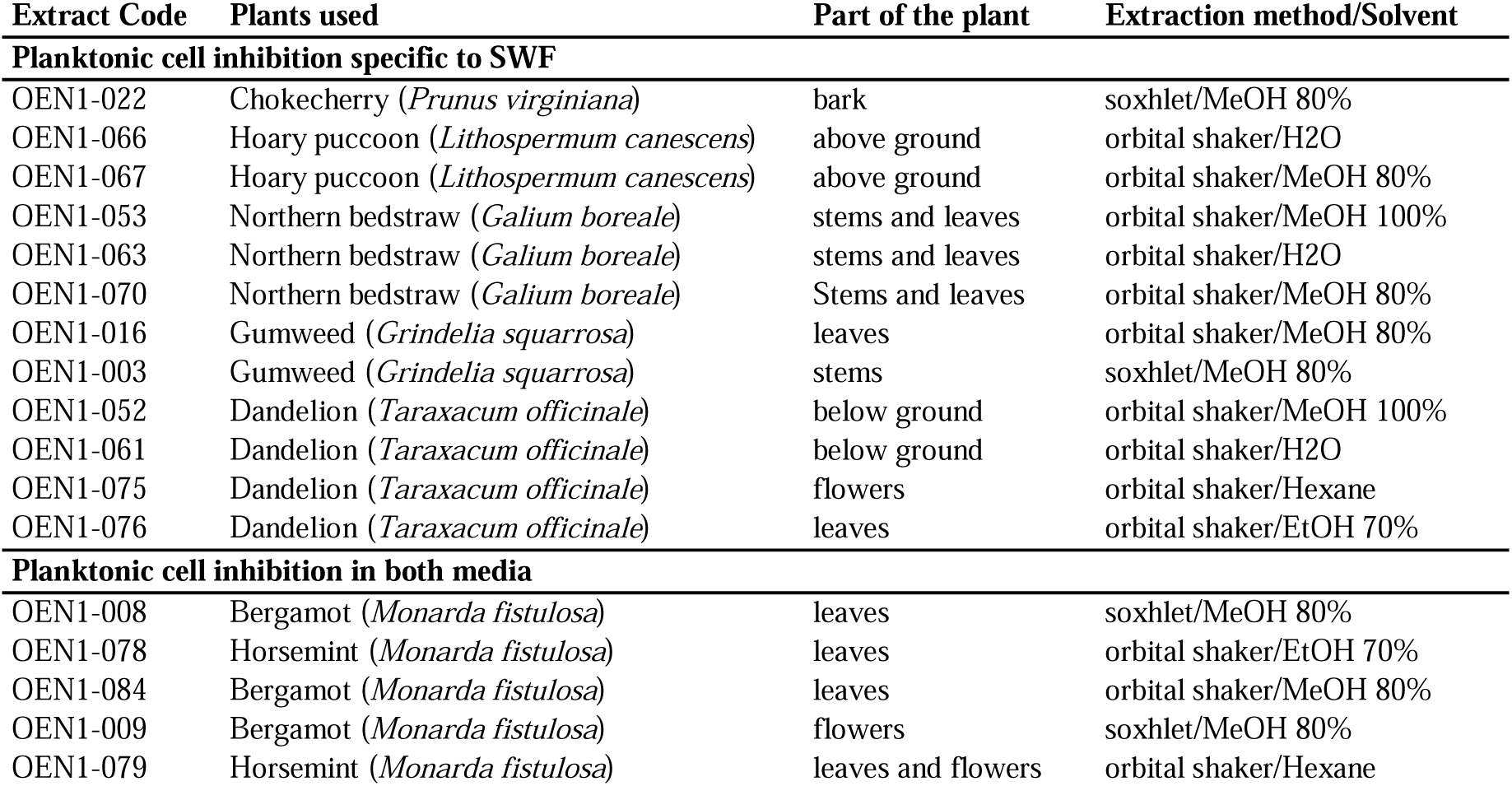
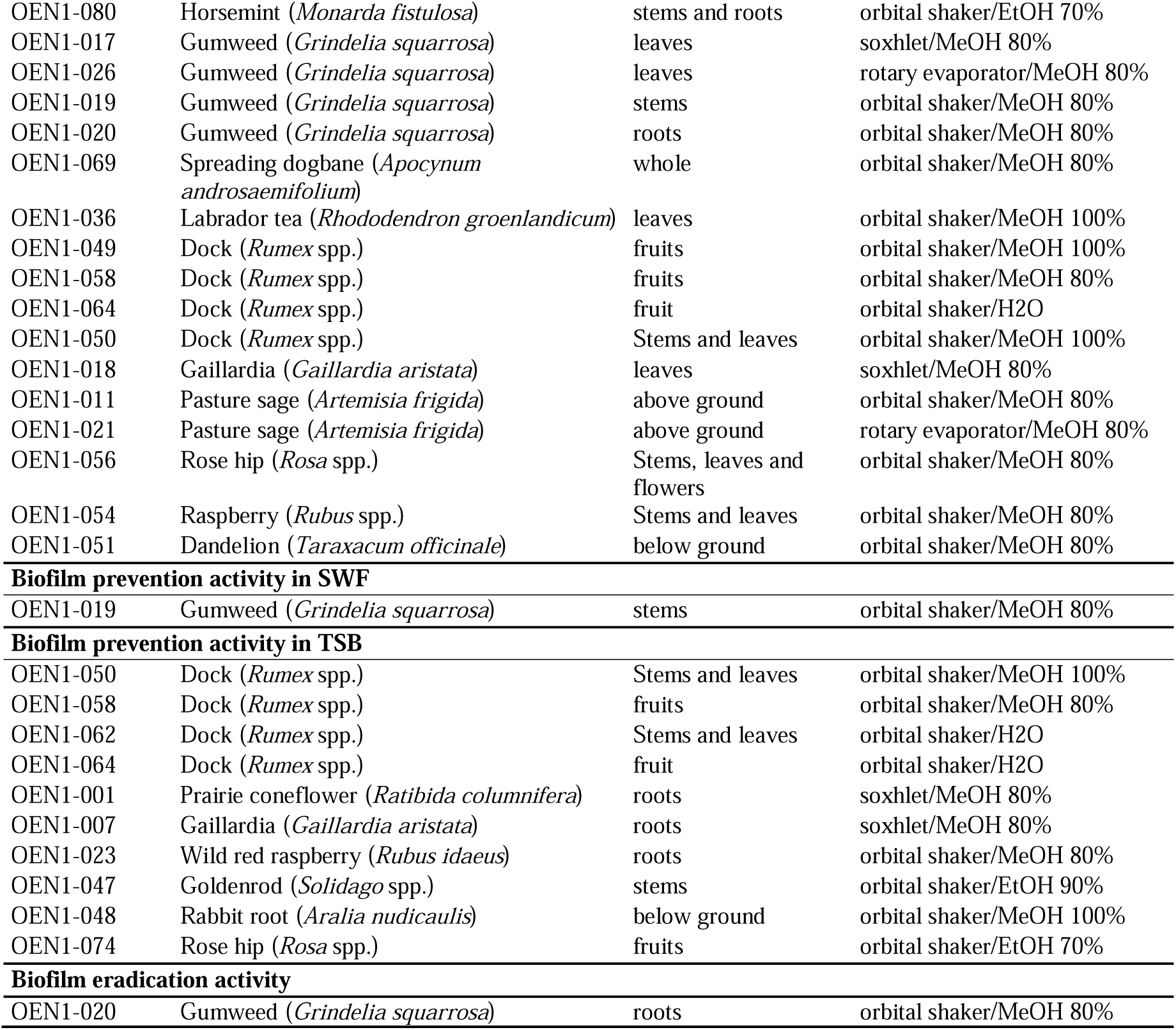
Select Indigenous remedies’ extracts used in this study showing antimicrobial activity.

| Extract Code | Plants used | Part of the plant | Extraction method/Solvent |
| --- | --- | --- | --- |
| **Planktonic cell inhibition specific to SWF** |  |  |  |
| OEN1-022 | Chokecherry (*Prunus virginiana*) | bark | soxhlet/MeOH 80% |
| OEN1-066 | Hoary puccoon (*Lithospermum canescens*) | above ground | orbital shaker/H<sub>2</sub>O |
| OEN1-067 | Hoary puccoon (*Lithospermum canescens*) | above ground | orbital shaker/MeOH 80% |
| OEN1-053 | Northern bedstraw (*Galium boreale*) | stems and leaves | orbital shaker/MeOH 100% |
| OEN1-063 | Northern bedstraw (*Galium boreale*) | stems and leaves | orbital shaker/H<sub>2</sub>O |
| OEN1-070 | Northern bedstraw (*Galium boreale*) | Stems and leaves | orbital shaker/MeOH 80% |
| OEN1-016 | Gumweed (*Grindelia squarrosa*) | leaves | orbital shaker/MeOH 80% |
| OEN1-003 | Gumweed (*Grindelia squarrosa*) | stems | soxhlet/MeOH 80% |
| OEN1-052 | Dandelion (*Taraxacum officinale*) | below ground | orbital shaker/MeOH 100% |
| OEN1-061 | Dandelion (*Taraxacum officinale*) | below ground | orbital shaker/H<sub>2</sub>O |
| OEN1-075 | Dandelion (*Taraxacum officinale*) | flowers | orbital shaker/Hexane |
| OEN1-076 | Dandelion (*Taraxacum officinale*) | leaves | orbital shaker/EtOH 70% |
| **Planktonic cell inhibition in both media** |  |  |  |
| OEN1-008 | Bergamot (*Monarda fistulosa*) | leaves | soxhlet/MeOH 80% |
| OEN1-078 | Horsemint (*Monarda fistulosa*) | leaves | orbital shaker/EtOH 70% |
| OEN1-084 | Bergamot (*Monarda fistulosa*) | leaves | orbital shaker/MeOH 80% |
| OEN1-009 | Bergamot (*Monarda fistulosa*) | flowers | soxhlet/MeOH 80% |
| OEN1-079 | Horsemint (*Monarda fistulosa*) | leaves and flowers | orbital shaker/Hexane |
| OEN1-080 | Horsemint (*Monarda fistulosa*) | stems and roots | orbital shaker/EtOH 70% |
| OEN1-017 | Gumweed (*Grindelia squarrosa*) | leaves | soxhlet/MeOH 80% |
| OEN1-026 | Gumweed (*Grindelia squarrosa*) | leaves | rotary evaporator/MeOH 80% |
| OEN1-019 | Gumweed (*Grindelia squarrosa*) | stems | orbital shaker/MeOH 80% |
| OEN1-020 | Gumweed (*Grindelia squarrosa*) | roots | orbital shaker/MeOH 80% |
| OEN1-069 | Spreading dogbane (*Apocynum androsaemifolium*) | whole | orbital shaker/MeOH 80% |
| OEN1-036 | Labrador tea (*Rhododendron groenlandicum*) | leaves | orbital shaker/MeOH 100% |
| OEN1-049 | Dock (*Rumex* spp.) | fruits | orbital shaker/MeOH 100% |
| OEN1-058 | Dock (*Rumex* spp.) | fruits | orbital shaker/MeOH 80% |
| OEN1-064 | Dock (*Rumex* spp.) | fruit | orbital shaker/H2O |
| OEN1-050 | Dock (*Rumex* spp.) | Stems and leaves | orbital shaker/MeOH 100% |
| OEN1-018 | Gaillardia (*Gaillardia aristata*) | leaves | soxhlet/MeOH 80% |
| OEN1-011 | Pasture sage (*Artemisia frigida*) | above ground | orbital shaker/MeOH 80% |
| OEN1-021 | Pasture sage (*Artemisia frigida*) | above ground | rotary evaporator/MeOH 80% |
| OEN1-056 | Rose hip (*Rosa* spp.) | Stems, leaves and flowers | orbital shaker/MeOH 80% |
| OEN1-054 | Raspberry (*Rubus* spp.) | Stems and leaves | orbital shaker/MeOH 80% |
| OEN1-051 | Dandelion (*Taraxacum officinale*) | below ground | orbital shaker/MeOH 80% |
| **Biofilm prevention activity in SWF** |  |  |  |
| OEN1-019 | Gumweed (*Grindelia squarrosa*) | stems | orbital shaker/MeOH 80% |
| **Biofilm prevention activity in TSB** |  |  |  |
| OEN1-050 | Dock (*Rumex* spp.) | Stems and leaves | orbital shaker/MeOH 100% |
| OEN1-058 | Dock (*Rumex* spp.) | fruits | orbital shaker/MeOH 80% |
| OEN1-062 | Dock (*Rumex* spp.) | Stems and leaves | orbital shaker/H2O |
| OEN1-064 | Dock (*Rumex* spp.) | fruit | orbital shaker/H2O |
| OEN1-001 | Prairie coneflower (*Ratibida columnifera*) | roots | soxhlet/MeOH 80% |
| OEN1-007 | Gaillardia (*Gaillardia aristata*) | roots | soxhlet/MeOH 80% |
| OEN1-023 | Wild red raspberry (*Rubus idaeus*) | roots | orbital shaker/MeOH 80% |
| OEN1-047 | Goldenrod (*Solidago* spp.) | stems | orbital shaker/EtOH 90% |
| OEN1-048 | Rabbit root (*Aralia nudicaulis*) | below ground | orbital shaker/MeOH 100% |
| OEN1-074 | Rose hip (*Rosa* spp.) | fruits | orbital shaker/EtOH 70% |
| **Biofilm eradication activity** |  |  |  |
| OEN1-020 | Gumweed (*Grindelia squarrosa*) | roots | orbital shaker/MeOH 80% |

### Growth inhibitory activity of Indigenous remedies’ extracts against planktonic bacteria

We conducted a screen of 85 Indigenous remedies’ plant extracts at 2% v/v (of the extract stock solution) for inhibitory effects of *S. aureus* USA300, a hypervirulent community-associated MRSA in MHB and synthetic wound fluid (SWF). A total of 34 extracts inhibited growth below 20% in either MHB or SWF (Figure 1A). Dose-response assays revealed that 33 of these extracts showed antimicrobial activity, either completely or partially inhibiting bacterial growth (Figure 1B), where 12 extracts from five plants [chokecherry (*Prunus virginiana*; Figure 1C), hoary puccoon (*Lithospermum canescens*; Figure 1D), Northern bedstraw (*Galium boreale*; Figure 1E), gumweed (*Grindelia squarrosa*; Figure 1F) and dandelion (*Taraxacum officinale;* Figure 1G)] were more active in SWF and 21 showed activity in both SWF and MHB (Figure S1).

**Figure 1.**
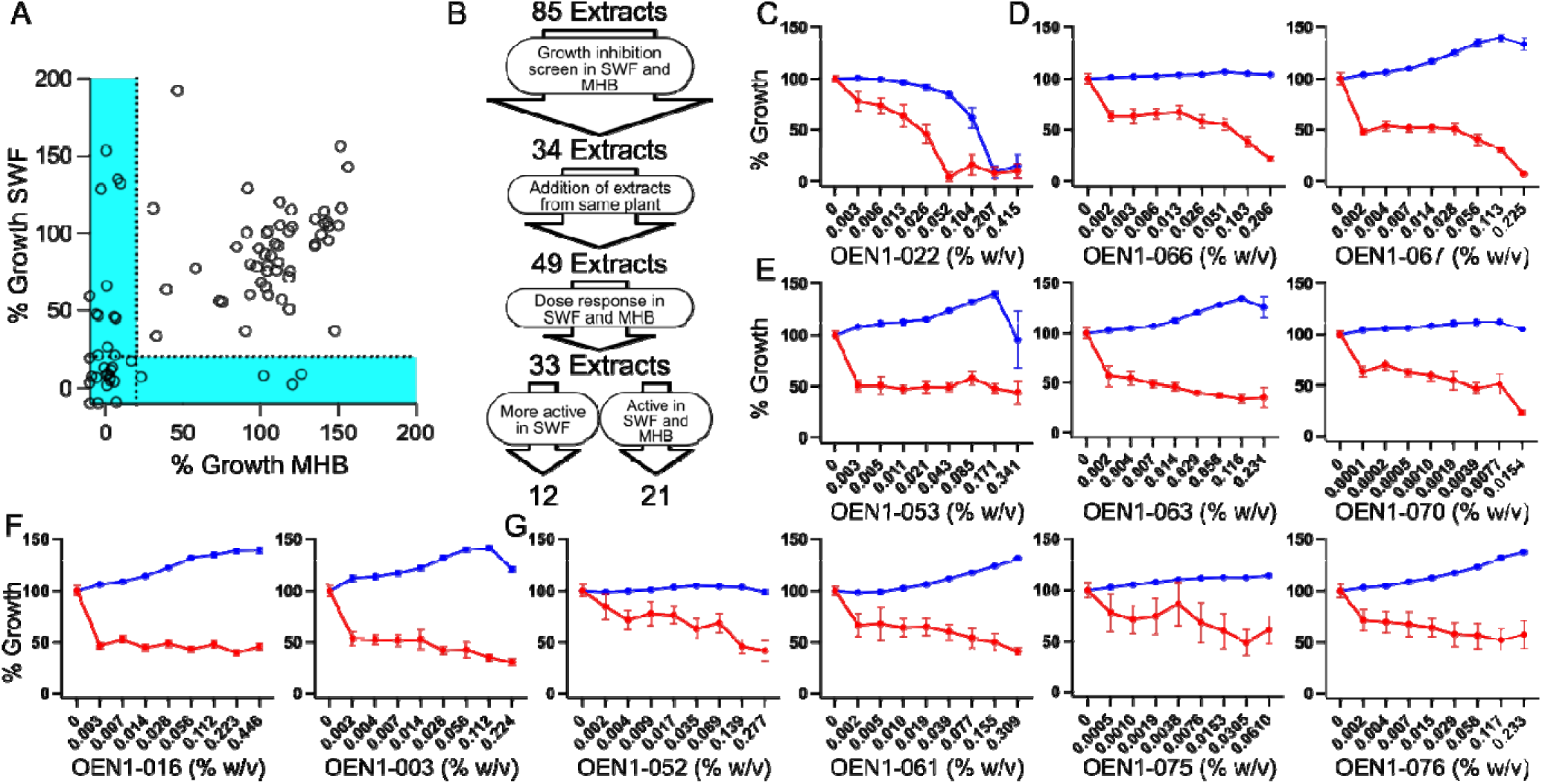
Screening of Indigenous remedies’ extracts for growth inhibitory activity against planktonic MRSA bacteria. A) Percent growth of MRSA USA300 treated with OEN1-001 to OEN1-085 relative to an untreated control cultured in SWF plotted against that in MHB. B) A flowchart of the growth inhibitory screen and dose-response assays that identified extracts more active in SWF or active in SWF and MHB. Dose-response curves of extracts of chokecherry (C), hoary puccoon (D), Northern bedstraw (E), gumweed (F), and dandelion (G), showing higher activity in SWF (red) relative to that in MHB (blue). N=6 from 3 independent experiments shown as mean of percent growth ± SEM.

### Indigenous remedies’ extracts with more potent antimicrobial activity in wound-mimetic conditions

The extract of the chokecherry bark (OEN1-022) inhibited the growth of USA300 in SWF at ∼4-fold lower concentration than in MHB, with minimum inhibitory concentration (MIC) values of 0.052% w/v compared to 0.207% w/v, respectively (Figure 1C). Two extracts of above ground parts of hoary puccoon, OEN1-066 and −067, showed gradual inhibition of USA300 growth over a wide concentration range leading to ∼80 and 100% inhibition, respectively, at ∼0.2% w/v (the highest tested concentration) only in SWF with no effects in MHB within the tested concentration range (Figure 1D). Three extracts from the stems and leaves of Northern bedstraw, OEN1-053, −063, and −070, also showed growth inhibitory effects in SWF only in the dose-response assays against USA300 (Figure 1E). The three extracts resulted in partial inhibition over a wide concentration range without completely inhibiting growth within the tested concentration range; OEN1-053, −063, and −070 showed MIC_50_ values (minimum concentration required to inhibit 50% of bacteria) of 0.171, 0.007, and 0.004% w/v, respectively (Figure 1E). The three Northern bedstraw extracts did not show growth inhibition in MHB (Figure 1E). Similarly, two extracts from gumweed leaves (OEN1-016) and stems (OEN1-003) showed gradual growth inhibition in SWF but not in MHB across the tested concentration range (Figure 1F). Four extracts from dandelion below ground portions (OEN1-052 and −061), flowers (OEN1-075), and leaves (OEN1-076) also partially inhibited growth in SWF with no observed growth inhibition in MHB (Figure 1G). Interestingly, these results (that revealed antimicrobial activity in SWF but not MHB) partly agree with a previous report that could not detect bacterial growth inhibition by extracts of aerial portions (including leaves, flowers, stems, and fruits) from Northern bedstraw, gumweed, and dandelions when tested in standard media conditions (36). Together, these results reveal the promising antimicrobial potential of some Indigenous natural plants while providing a proof-of-concept for screening for antimicrobial activity in infection-mimetic conditions.

### Indigenous remedies’ extracts with antimicrobial activity in both standard and wound-mimetic media conditions

Extracts from bergamot, also known as horsemint (*Monarda fistulosa*) leaves (OEN1-008, −078, and −084), flowers (OEN1-009), and leaves and flowers (OEN1-079) showed growth inhibitory effects against USA300 in both MHB and SWF (Figure S1A). Some of these extracts showed comparable activity in both media (such as OEN1-084), while others showed reduced growth inhibition in SWF with OEN1-008 showing the largest MIC shift compared to MHB (∼64-fold, Figure S1A). Altered activity in one or both media across the different extracts from the same plant part might be attributed to differences in extraction procedures possibly concentrating varied chemical entities (Table 1). For example, OEN1-084 (MIC of 0.024 % w/v in MHB and 0.049 %w/v in SWF) and OEN1-008 (MIC of 0.004 %w/v in MHB and 0.142 %w/v in SWF) are 80% methanolic extracts of bergamot leaves prepared using an orbital shaker (OEN1-084) and a Soxhlet apparatus (OEN1-008). Interestingly, an extract of the stems and roots of bergamot (OEN1-080) did not show growth inhibitory effects under the test conditions (Figure S1A), suggesting the lack of active antimicrobial components in these plant parts and highlighting the importance of using the correct plant parts.

Extracts from gumweed leaves (OEN1-017 and OEN1-026), stems (OEN1-019), and roots (OEN1-020) also showed growth inhibitory effects against USA300 in both MHB and SWF (Figure S1B). Notably, extracts of the leaves (OEN1-016) and stems (OEN1-003) showed reduced antimicrobial activity in MHB compared to the other extracts of the same parts prepared by different methods in the same solvent (Table 1 and Figure 1F).

Extracts from spreading dogbane (*Apocynum androsaemifolium*) whole plant (OEN1-069; Figure S1C), Labrador tea (*Rhododendron groenlandicum*) leaves (OEN1-036; Figure S1D), dock (*Rumex* spp.) fruit (OEN1-049, −058, and −064; Figure S1E) and stems and leaves (OEN1-050; Figure S1E), and gaillardia (*Gaillardia aristata*) leaves (OEN1-018; Figure S1F) also showed growth inhibitory effects against USA300 in both MHB and SWF with comparable activities in both media. Extracts of other plants, pasture sage (*Artemisia frigida*) (OEN1-011 and −021; Figure S1G), rose hip (*Rosa* spp.) (OEN1-056; Figure S1H), wild raspberry (*Rubus* spp.) (OEN1-054; Figure S1I), and dandelion (OEN1-051; Figure S1J), also resulted in growth reduction in both media, reaching complete growth inhibition in both media (by OEN1-011 and −056) and MHB only (by OEN1-021, −054, and - 051).

### Cross-resistance platform screen de-replicates the bacterial growth inhibitory Indigenous remedies’ natural product extracts

Natural product discovery is ridden with re-discovery and thus de-replication is an important step before progressing with natural product derived bioactives (37). An established de-replication strategy is to test newly identified bioactives against clones resistant to known antibiotics (38). As such, we sought to test 17 Indigenous remedies’ extracts that inhibited the growth of USA300 with higher potency in SWF (OEN1-022, −067, −053, −063, and −070; Figure 1) or with relatively comparable potency in SWF and MHB (OEN1-008, −078, −084, −079, −069, −036, −049, −058, −064, − 050, −011, and −021; Figure S1) against a *S. aureus* cross-resistance platform (CRP) that contains 23 strains encoding plasmids or mutations that confer resistance to most known classes of antibiotics in the *S. aureus* SH1000 parent strain (39) (Table S1). The hypothesis is that resistance of a CRP strain to an antimicrobial extract would suggest a similar mechanism of action of the extract and the antibiotic to which the CRP strain is resistant. First, we determined the MIC of the extracts against parent SH1000 strain in MHB and SWF (Table S2). Next, we tested the extracts that completely inhibited the growth of SH1000 within the tested concentration range (Table S2) against the CRP strains in each culture media at twice their respective SH1000 MIC in each medium (Figure 2A and B). Extracts OEN1-021 and −070 did not show an MIC against SH1000 in both culture media (Table S2), hence they were not tested against the CRP strains. Each CRP strain was resistant to its respective antibiotic control with at least 4-fold increase in the MIC relative to the parent strain in SWF and MHB (Table S1); Figure 2A and B shows the lack of growth inhibition of each strain by its antibiotic control at 2x the wild-type MIC in both media, validating the use of the CRP strains. Most extracts inhibited the growth of the CRP strains in both media (Figure 2A and B). Two extracts, OEN1-022 and OEN1-064, failed to completely inhibit the growth of certain CRP strains in MHB (Figure 2B). However, follow-up dose-response assays of OEN1-022 (against AJUL 5 and 22, Figure 2C), and OEN1-064 (against AJUL 8, 16, and 24, Figure 2D) in SWF and MHB revealed no notable difference in the MIC against these CRP strains and their parent SH1000 strain. Together, these results suggest novel antimicrobial mechanisms of action, different from those of currently known antibiotics, for the extracts of the Indigenous remedies described in this study, paving the way for future studies to characterize their active antimicrobial components and their mode of action.

**Figure 2.**
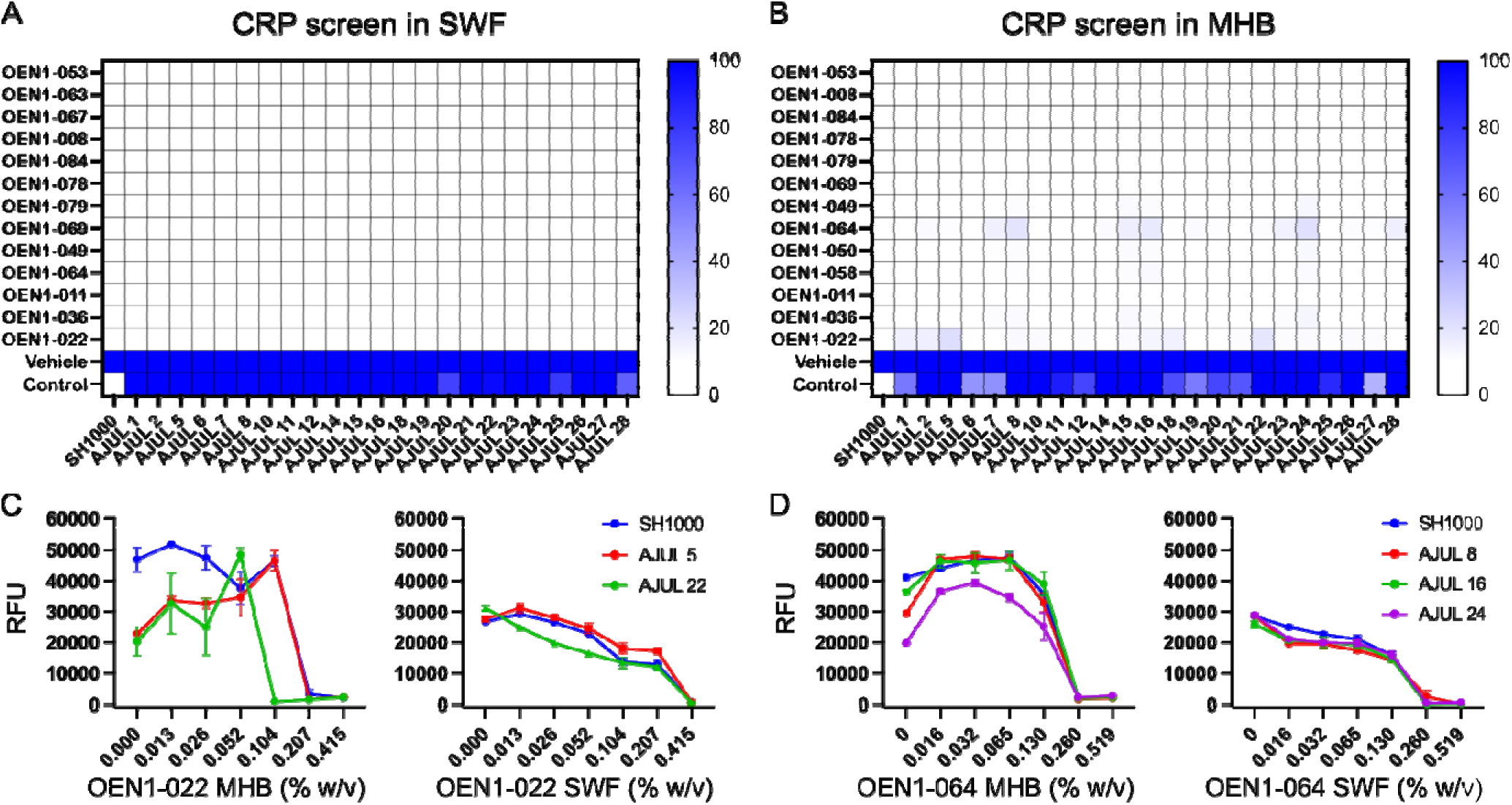
Cross-resistance platform (CRP) screen to de-replicate the bacterial growth inhibitory natural product extracts. A and B) Heat map of the growth of CRP strains and its parent SH1000 strain in the presence of the extracts at 2X the MIC of SH1000 in SWF (A) and MHB (B) shown as percentage of their growth in the presence of the vehicle only (N=1). The vehicle is 4 µL of DMSO while the control is 2x the MIC of SH1000 of the respective antibiotic control for each CRP strain (Table S1). C and D) Dose-response curves of OEN1-022 (C) and OEN1-064 (D) against SH1000 and certain CRP strains that showed growth in the CRP screen (N=2 shown as mean ± SEM).

### Biofilm prevention activity of Indigenous remedies’ extracts

We screened the 85 Indigenous remedies’ plant extracts at 2% v/v for MRSA USA300 biofilm prevention in SWF compared to the standard medium TSB, both in the presence of 1% glucose. We identified 37 extracts with ≥60% reduction in biofilm formation relative to the untreated control detected as the absorbance signal of the solubilized crystal violet dye staining the biofilm biomass (Figure 3A). Next, we excluded the extracts showing planktonic growth inhibition, yielding 20 extracts that showed biofilm prevention but not growth inhibitory activity (Figure 3B); we tested these extracts in subsequent dose-response assays (Figure 3C, and Figure S2). The dose-response results identified one extract (OEN1-019) that showed activity in SWF only (Figure 3 and Table 1). OEN1-019, extracted from the stems of gumweed, showed ∼58% biofilm prevention activity in SWF at its highest tested concentration (0.215% w/v) compared to the untreated control (Figure 3C). Notably, another ten extracts from seven plants showed biofilm prevention activity in TSB (Figure S2 and Table 1). Four extracts (OEN1-050, −058, −062, and −064) were extracted from the stems, leaves and fruits of dock plant. The other six extracts (OEN1-001, −007, −023, −047, −048, and −074) were from prairie coneflower (*Ratibida columnifera*), gaillardia, wild red raspberry (*Rubus idaeus*), goldenrod (*Solidago* spp.), rabbit root (*Aralia nudicaulis*), and rose hip, respectively (Table 1). All ten extracts showed more than 50% (up to complete) biofilm prevention in a dose dependant manner; the most potent of which was OEN1-023, showing total biofilm prevention (91%) at 0.028% w/v followed by OEN1-007 showing 86.6% biofilm prevention at 0.029% w/v (Figure S2).

**Fig 3.**
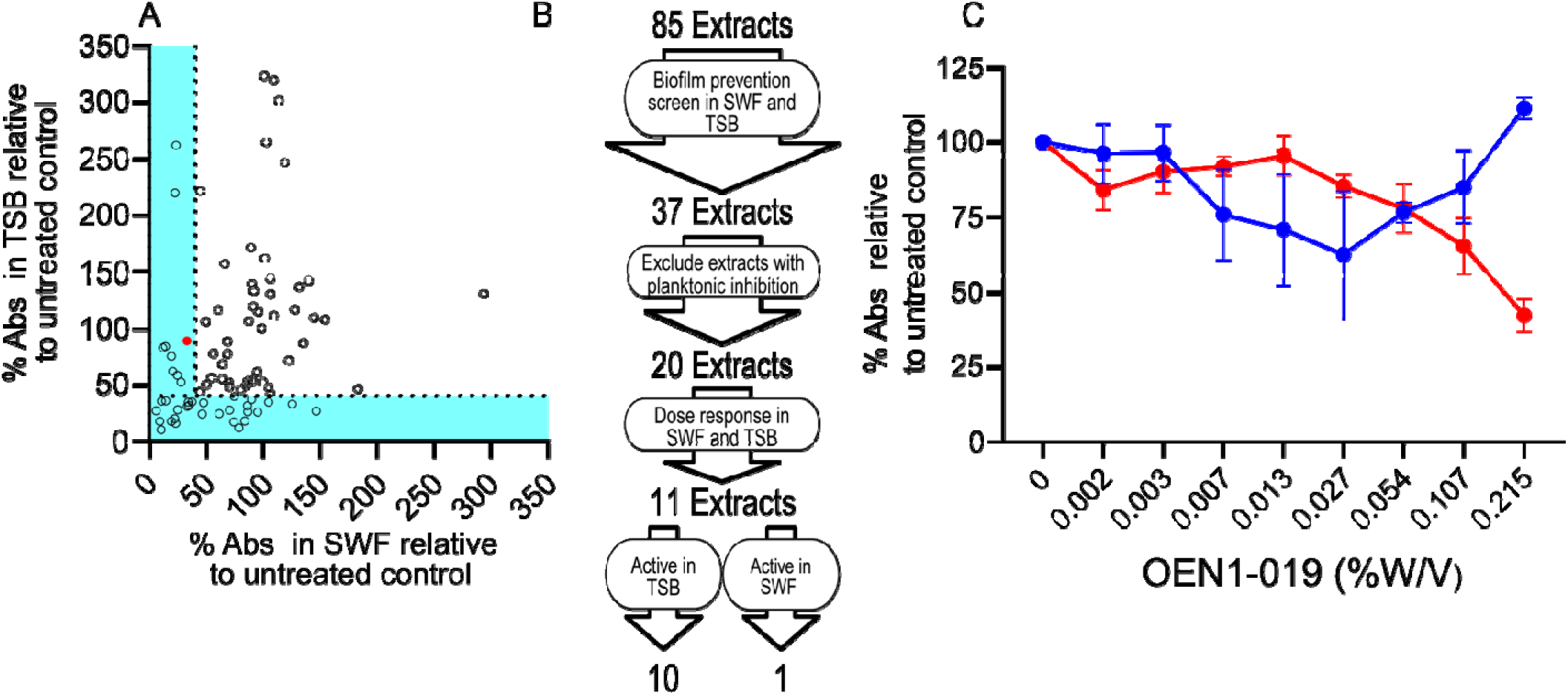
Screening of Indigenous remedies’ extracts for biofilm prevention activity against MRSA. A) Percent A_590_ of USA300 biofilm treated with OEN1-001 to OEN1-085 relative to an untreated control cultured in SWF plotted against that in TSB. B) A flowchart of the biofilm prevention screen and dose-response assays that identified extracts active in SWF (red) or in TSB (blue). C) Dose-response assay of OEN1-019 showing ∼ 58% biofilm prevention activity at 0.215 % w/v in SWF relative to the untreated control. The red colored small circle in (A) indicates OEN1-019. N=4 from 2 independent experiments shown as mean percent absorbance ± SEM.

### Biofilm eradication activity of Indigenous remedies’ extracts

We also screened the Indigenous remedies’ plant extracts at 4% v/v for the eradication of pre-formed MRSA USA300 biofilms in SWF compared to the standard medium TSB, both in the presence of 1% glucose (Figure 4A). We considered 36 that eradicated ≥40% of the pre-formed biofilm relative to the untreated control detected as the crystal violet absorbance signal (Figure 4B) for follow up dose-response assays (Figure 4C and Figure S3). These dose-response assays revealed that only one plant extract (OEN1-020) possessed a dose-dependent biofilm eradication activity in both SWF and TSB with maximal activity observed at 0.189% w/v (Figure 4C) above which this activity was gradually lost either due to reduced effective concentration resulting from potential extract precipitation at higher concentrations or partial biofilm enhancement. OEN1-020 is an 80% methanolic extract of gumweed roots prepared using the orbital shaker (Table 1).

**Figure 4.**
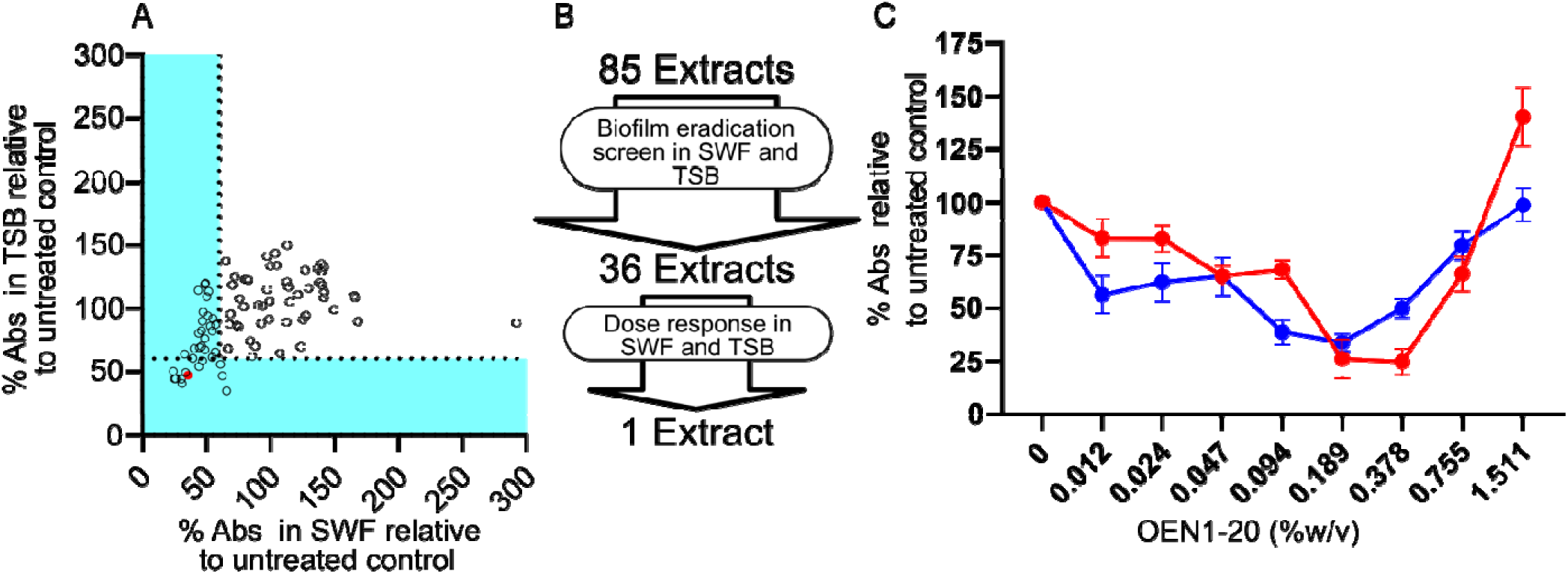
Screening of Indigenous remedies’ extracts for biofilm eradication activity against MRSA. A) percent A_590_ of USA300 preformed biofilm treated with OEN1-001 to OEN1-085 relative to an untreated control cultured in SWF plotted against that in TSB. B) A flowchart of the biofilm eradication screen and dose-response assays that identified one extract active in both media. C) Dose-response assay of OEN1-020 showing ∼ 84% and ∼ 66% biofilm eradication activity at 0.189 % w/v in SWF (red) and TSB (blue), respectively, relative to the untreated control. The red colored small circle in (A) indicates OEN1-020. N=8 from 3 independent experiments shown as mean percent absorbance ± SEM.

Interestingly, a total of nine extracts in the biofilm prevention screen (eight in TSB and one in SWF; Figure 3A) and one extract in the biofilm eradication screen in SWF (OEN1-063; Figure 4A) stimulated the biofilm production shown as more than double the crystal violet absorbance signals relative to the untreated control. These extracts demonstrated some growth inhibitory activity against planktonic cells in the same media, thus their biofilm stimulatory activity is consistent with previous reports of similar enhancement of biofilm production by antibiotics at sub-inhibitory levels (40, 41). For example, low level β-lactam exposure stimulated biofilm production in *S. aureus* by releasing extracellular DNA (40).

## Conclusions

Here, we tested the antimicrobial properties of an array of Indigenous remedies’ extracts against MRSA under infection-mimetic conditions. Unlike typical antibiotic discovery campaigns, we combined high-throughput and novel drug discovery approaches with exploring traditional Indigenous medicines and knowledge. Our goal was to co-create new antimicrobial solutions for drug-resistant MRSA infections inspired by Indigenous medicines and forge a two-way stream for knowledge dissemination and translation between Indigenous and non-Indigenous scientists to address the MRSA crisis that constitutes an Indigenous and non-Indigenous health priority. Our work may also provide a roadmap that inspires similar interdisciplinary collaborations between Indigenous and non-Indigenous researchers to tackle other healthcare priorities and diseases.

Our choice to test the Traditional Indigenous remedies’ extracts against MRSA under wound infection-mimetic conditions was driven by our commitment to the principles of reciprocity and mutual benefit when exploring Indigenous knowledge and partnering with Indigenous People (42). MRSA wound infections constitute an Indigenous health priority (7, 9–11); thus promising positive results from our collaborative work may be amenable to a relatively rapid implementation within Indigenous communities should Indigenous Elders wish to repurpose such Traditional remedy for topical administration to fight wound infection.

The antimicrobial potential of some of the extracts tested herein warrants further studies to identify the active components in each plant extracts and explore the mechanism of their antimicrobial activity. However, any such future steps also warrant conversations regarding data sovereignty, intellectual property, potential commercialization prospects, and guarding against cultural appropriation. Such discussions are necessary to address any legitimate concerns mediated by long repeating breaches of trust incurred against Indigenous populations in Canada and worldwide by non-Indigenous peoples (6). Indeed, such conversations will be beneficial in the areas of antibiotic drug discovery and development but also for other areas of drug discovery should this interdisciplinary approach expand to other human disease applications like cancer.

Taken together, we assembled a natural product screening library comprised of extracts of an array of plants native to the Canadian Prairies that are of medicinal value for Indigenous Peoples. We undertook an interdisciplinary partnership that leverages Western approaches and traditional Indigenous knowledge to co-create new antimicrobial solutions. We identified plant extracts with promising antibacterial growth inhibitory activity [extracts from chokecherry, hoary puccoon, Northern bedstraw, gumweed, dandelion, bergamot/horsemint, spreading dogbane, Labrador tea, dock, gaillardia, pasture sage, rose hip, and raspberry], biofilm prevention [extract from gumweed], and biofilm eradication [extract from gumweed] against MRSA under wound infection-mimetic conditions (summarized in Table 1). These extracts may offer new solutions to combat drug resistant MRSA wound infections.

## Materials and Methods

### Plant collection

All plant collection was carried out with the assistance of Indigenous Elders, Medicine People, and Traditional Knowledge Keepers throughout three ecozones and several communities in Treaties 4, 6, and 10 in Saskatchewan, Canada. Relationship building between researchers, students, and the Elders participating was necessary before educational Medicine Walks or picking sessions occurred. First, Tobacco in the form of a Tobacco Tie (bundle of Tobacco) was offered to each Elder while communicating the intended purpose of the research and request for sharing of Traditional Knowledge about Medicinal Plant Healing Traditions. Each harvesting session followed a Tobacco offering and commenced after the provision of Honouraria (monetary offering of thanks for time and Knowledge). In most cases, a small amount of loose Tobacco was offered at the site of harvest with the assistance of the Traditional Knowledge Keeper before picking the plants. Great care was taken to remove the whole plant without damage when feasible, and only a fraction of the plant species in the area was harvested to ensure health and good yields during the next harvesting period for each species.

### Indigenous remedies extraction

The extraction of whole plants or plant parts, such as leaves, stems, roots, flowers, bark, fruits or a combination thereof was performed by orbital shaker, Soxhlet, rotary evaporator, or sonication in water, methanol (ranging from 100% to 70% diluted with water), ethanol (ranging from 90% to 70% diluted with water), or hexane. The method and solvent used for extracts of interest are specified in Table 1. After extraction, all extracts were dried under vacuum using DNA 110 SpeedVac® concentrator. Concentrator cycles consisted of 1h at room temperature under vacuum followed by 3h at 45 °C under vacuum. Cycles were repeated until the extracts were completely dried when the final dry weight was recorded. All dried extracts were then re-dissolved in 1.00 mL of 99.7% DMSO by vortexing. Extract stock concentrations varied from 0.7 mg/mL to 637.2 mg/mL and aliquots were stored at −80°C.

### Strains and reagents

We used the *S. aureus* USA300 LAC, which is a hypervirulent community-associated MRSA isolated in 2002 from a skin and soft tissue infection of an inmate in the Los Angeles County Jail in California, USA., cured of its plasmids (also known as JE2) for all experiments unless otherwise indicated. We also used the *S. aureus* cross-resistance platform (CRP) that contains 23 strains encoding plasmids or mutations that confer resistance to most known classes of antibiotics in the *S. aureus* SH1000 parent strain (39). Bacteria were grown in cation-adjusted Mueller Hinton broth (MHB), tryptic soy broth supplemented with 1% glucose (TSB), or synthetic wound fluid composed of 50% fetal bovine serum (Fisher Scientific and VWR) and 50% maximum recovery diluent (MRD; 1g/L peptone and 8.5g/L NaOH). Glucose was added at 1% to SWF for biofilm assays. Bacteria were incubated at 37 °C.

### Growth inhibitory screen

Two microliters of plant extracts were added to 98 µL of either MHB or SWF dispensed into each well in a microtiter plate. USA300 was inoculated using the colony resuspension method (43) to a final inoculum size at an OD_600_ of 0.001. Plates were incubated for 24 h at 37°C with shaking. Growth values were background subtracted by wells containing uninoculated media with 2 µL of the plant extract. Percent growth was determined by dividing the background subtracted value by the vehicle control.

### Antimicrobial Susceptibility Testing

Minimum inhibitory concentrations (MIC) were determined following CLSI guidelines for MIC testing by broth microdilution(43). Reading of plates followed the same method as described in growth inhibitory screen.

### Cross-resistance screen

Four microliters of plant extracts were dispensed into microtiter plate wells containing 96 µL of MHB or SWF; the final extract concentration was equivalent to 2x MIC against the wild-type strain. Plates were incubated at 37°C for 24 h followed by the addition of 5 µL of 0.15 mg/mL resazurin. The plates were further incubated for two hours in the dark and read at 554nm/593nm excitation/emission wavelengths.

### Biofilm inhibition assay

The biofilm inhibition activity of the indigenous natural plant extracts library against USA300 was assessed using a crystal violet (CV) staining method as previously reported (44). Briefly, a bacterial suspension of USA300 was diluted to a final OD_600_ of 0.01 in TSB or SWF with 1% glucose. Then, 49 µL of each medium and 1 µL of each plant extract were added to a 384 wells polystyrene Nunc plates (for library screening). The plates were incubated under static conditions for 24h at 37 °C. OD_600_ of planktonic growth was recorded followed by discarding the cultures. Next, we washed the plates with deionized water, dispensed 0.1% CV, discarded the excess CV, washed the plates again with deionized water, and dispensed 30% acetic acid using the BioTek EL406 microplate washer/dispenser (Agilent, Santa Clara, CA, USA). The absorbance of CV-stained biofilms was recorded at 590 nm. Then, the results were documented as % Abs relative to the untreated control. The same method was used in the dose response by mixing 2 µL of each plant extract with 98 µL of each medium in 96 wells polystyrene Nunc plates.

### Biofilm eradication assay

The biofilm eradication screen was performed on a pre-established biofilms of USA300 in 96-well Nunc plates as previously reported (44). Briefly, 100 µL TSB or SWF plus 1% glucose inoculated with USA300 bacterial suspension of OD_600_ = 0.01 were added. After 24h of static incubation at 37 °C, the plates were washed with sterile deionized water using BioTek EL406 washer/dispenser. Then, 96 µL of each medium and 4 µL of each plant extract were added to different wells. After 24h of static incubation at 37 °C, the plates underwent CV staining assay as mentioned above. The results were documented as % Abs relative to the untreated control. For the dose response assays, the extracts were 2-fold serially diluted and added to the cultures as described above for biofilm prevention and eradication assays.

## Acknowledgements

This work was funded by a New Frontiers in Research Funds exploration grant (NFRFE-2021-00398). O.M.E. holds a Canada Research Chair in Chemogenomics and Antimicrobial Research (CRC-2019-00115). C.D.R. was supported by a Canadian Institutes of Health Research Canada Graduate Scholarship – Master’s (CGS-M) award. K.P., N.G., A.C.L., S.A.V., A.G.G.F., and M.A.F.B. were supported by Mitacs Globalink Research Internship awards. We thank Elders William Ratfoot (Makwa Sahgaiecan First Nation), Dennis Omeasoo (Maskwacis and Piapot First Nations), and Margaret Rocktuhunder (Piapot First Nation), as well as Oskâpêwis Roland Kaye (Sakimay First Nation) for their guidance.

