## Supplementary material for "The antimicrobial potential of traditional remedies of Indigenous Peoples from Canada against MRSA planktonic and biofilm bacteria in wound-infection mimetic conditions": Supp Figures and tables

[†]Deceased on 7 January 2023

[††]Deceased on 3 July 2022

[†††]Deceased on 2 September 2024


**Running title: The antimicrobial potential of Indigenous remedies against MRSA**

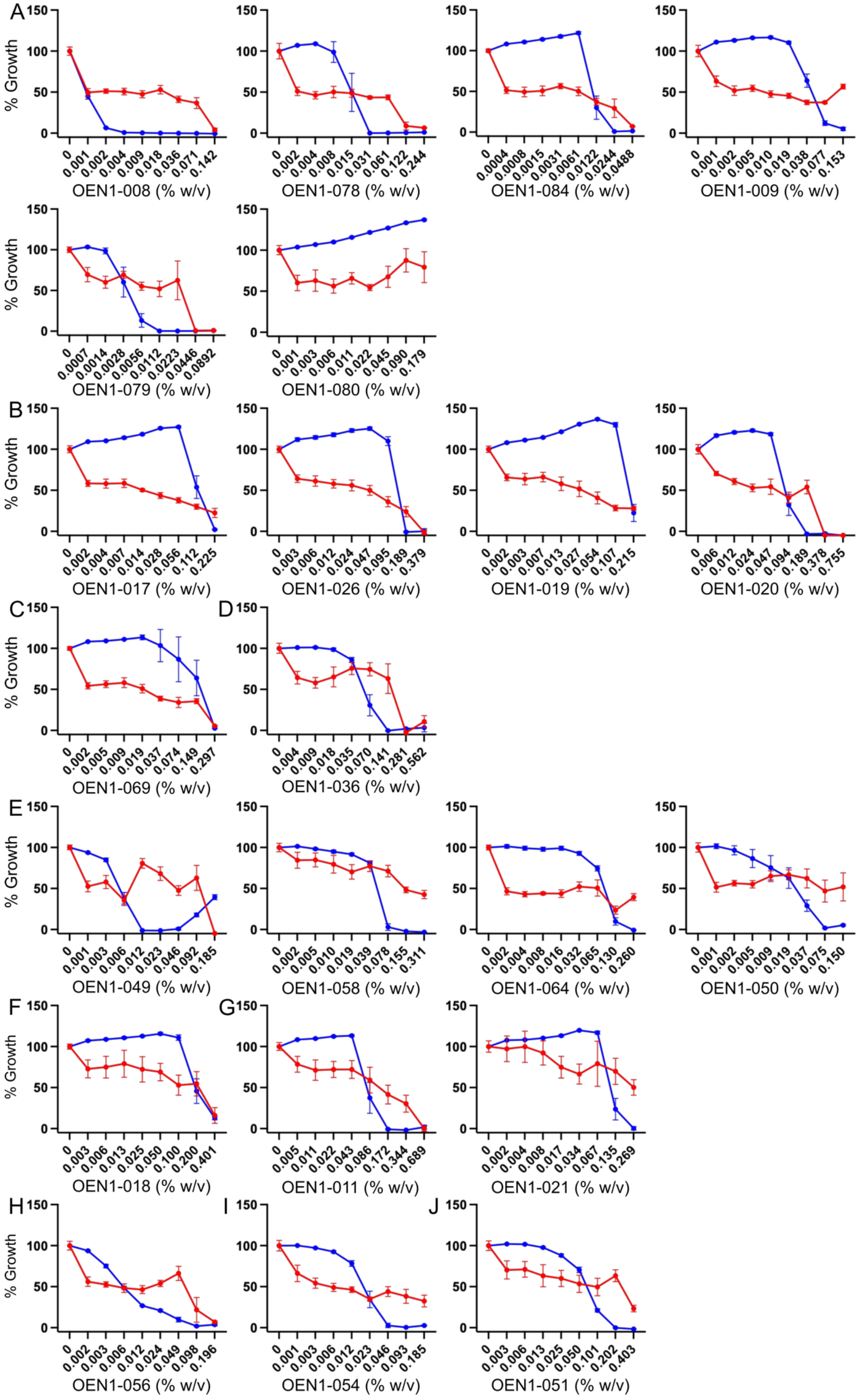
A
B
C
D
E
F
G
H
I
J
% Growth
150
100
50
0
0 0.001 0.002 0.004 0.009 0.018 0.036 0.071 0.142
OEN1-008 (% w/v)
0 0.002 0.004 0.008 0.015 0.031 0.061 0.122 0.244
OEN1-078 (% w/v)
0 0.0004 0.0008 0.0015 0.0031 0.0061 0.0122 0.0244 0.0488
OEN1-084 (% w/v)
0 0.001 0.002 0.005 0.010 0.019 0.038 0.077 0.153
OEN1-009 (% w/v)
0 0.0007 0.0014 0.0028 0.0056 0.0112 0.0223 0.0446 0.0892
OEN1-079 (% w/v)
0 0.001 0.003 0.006 0.011 0.022 0.045 0.090 0.179
OEN1-080 (% w/v)
0 0.002 0.004 0.007 0.014 0.028 0.056 0.112 0.225
OEN1-017 (% w/v)
0 0.003 0.006 0.012 0.024 0.047 0.095 0.189 0.379
OEN1-026 (% w/v)
0 0.002 0.003 0.007 0.013 0.027 0.054 0.107 0.215
OEN1-019 (% w/v)
0 0.006 0.012 0.024 0.047 0.094 0.189 0.378 0.755
OEN1-020 (% w/v)
0 0.002 0.005 0.009 0.019 0.037 0.074 0.149 0.297
OEN1-069 (% w/v)
0 0.004 0.009 0.018 0.035 0.070 0.141 0.281 0.562
OEN1-036 (% w/v)
0 0.001 0.003 0.006 0.012 0.023 0.046 0.092 0.185
OEN1-049 (% w/v)
0 0.002 0.005 0.010 0.019 0.039 0.078 0.155 0.311
OEN1-058 (% w/v)
0 0.002 0.004 0.008 0.016 0.032 0.065 0.130 0.260
OEN1-064 (% w/v)
0 0.001 0.002 0.005 0.009 0.019 0.037 0.075 0.150
OEN1-050 (% w/v)
0 0.003 0.006 0.013 0.025 0.050 0.100 0.200 0.401
OEN1-018 (% w/v)
0 0.005 0.011 0.022 0.043 0.086 0.172 0.344 0.689
OEN1-011 (% w/v)
0 0.002 0.004 0.008 0.017 0.034 0.067 0.135 0.269
OEN1-021 (% w/v)
0 0.002 0.003 0.006 0.012 0.024 0.049 0.098 0.196
OEN1-056 (% w/v)
0 0.001 0.003 0.006 0.012 0.023 0.046 0.093 0.185
OEN1-054 (% w/v)
0 0.003 0.006 0.013 0.025 0.050 0.101 0.202 0.403
OEN1-051 (% w/v)

**Figure S1. Dose-response assays of Indigenous remedies' extracts** from A) bergamot (horsemint) OEN1-008, -078, -084, -009, -079, and -080; B) gumweed OEN1-017, -026, -019, and -020; C) spreading dogbane OEN1-069; D) labrador tea OEN1-036; E) dock OEN1-049, -058, -064, and -050; F) gaillardia OEN1-018; G) pasture sage OEN1-011 and -021; H) rose hip OEN1-056; I) wild raspberry OEN1-054; J) dandelion OEN1-051. Dose-response curves in SWF (red) and MHB (blue). N=6 from 3 independent experiments shown as mean of the percent growth ± SEM.

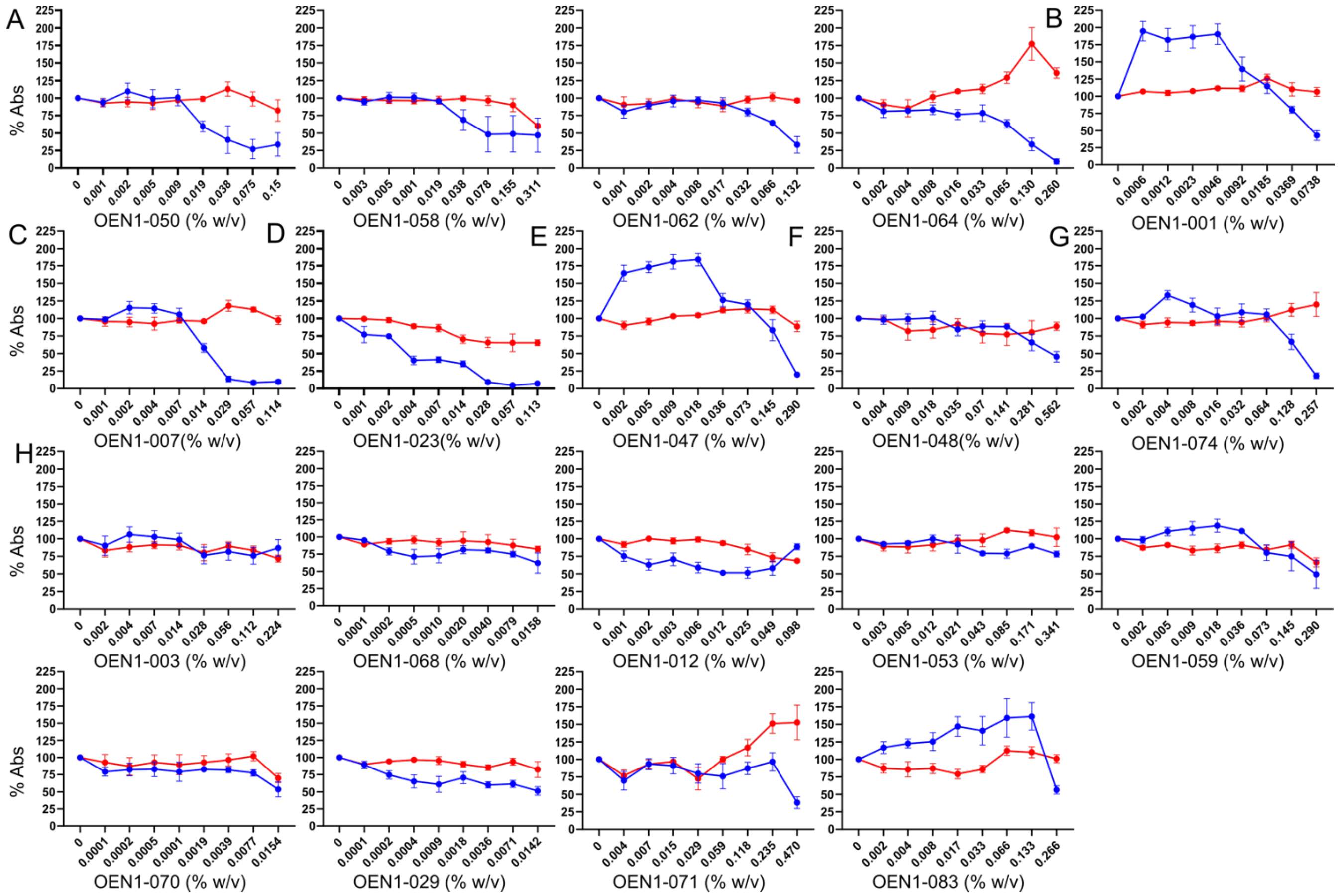


**Figure S2. Dose-response assays of Indigenous remedies' extracts for biofilm prevention activity against MRSA.** A) Dock OEN1-050, -058, -062, and -064; B) prairie coneflower OEN1-001; C) Gaillardia OEN1-007; D) wild red raspberry OEN1-023; E) goldenrod OEN1-047; F) rabbit root OEN1-048 G) rose hip OEN1-074; H) represent extracts without biofilm prevention activity in TSB or SWF. Y-axis represents percent $A_{590}$ of USA300 biofilm treated with the corresponding extract relative to an untreated control. Dose-response curves are in SWF (red) and TSB (blue). The results were shown as mean percent absorbance ± SEM.

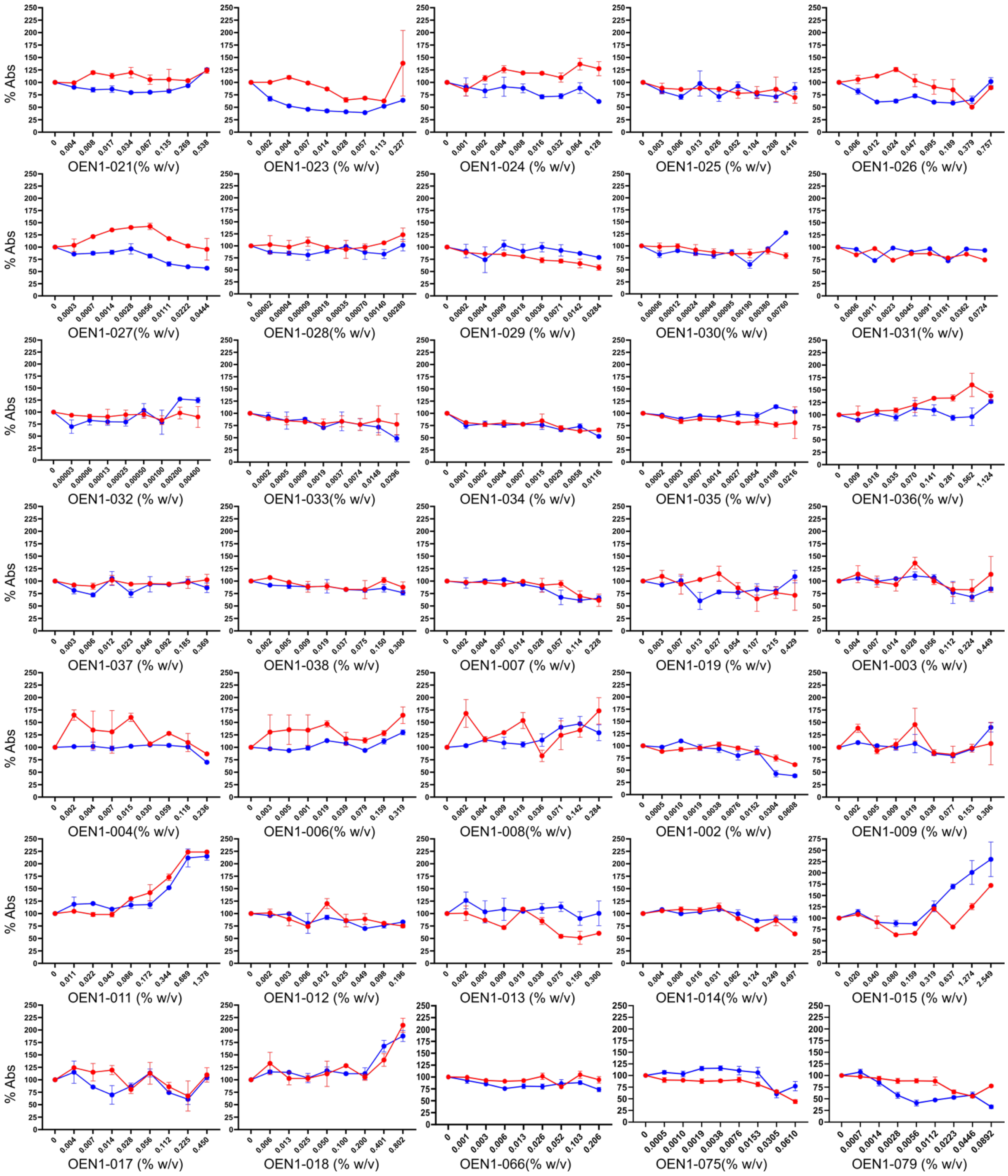


**Figure S3. Dose-response assays of Indigenous remedies' extracts for biofilm eradication activity against MRSA.** The figure represents extracts without notable activity either in TSB or in SWF. Y-axis represents percent $A_{590}$ of USA300 pre-formed biofilm treated with the corresponding extract relative to an untreated control. Dose-response curves are in SWF (red) and TSB (blue). The results were shown as mean percent absorbance ± SEM.

**Table S1.** MIC values (μg/mL) of control antibiotics tested against SH1000 and CRP strains cultured in SWF or MHB

| Strain tested | Control antibiotic tested | CRP strain | | SH1000 | | Resistant to antibiotic class |
|---|---|---|---|---|---|---|
| | | MHB | SWF | MHB | SWF | |
| AJUL1 | chloramphenicol | >64 | >64 | 8 | 8 | phenicols |
| AJUL2 | kanamycin | >16 | >16 | 0.5 | 0.5 | aminoglycosides |
| AJUL5 | kanamycin | >16 | >16 | 0.5 | 0.5 | aminoglycosides |
| AJUL6 | streptomycin | 512 | 32 | 16 | 1 | aminoglycosides |
| AJUL7 | retapamulin | >0.5 | >0.5 | 0.125 | 0.0625 | Phenicols, linxosamides, oxazolidinones, pleuromutilins, steptogramins (A) |
| AJUL8 | erythromycin | >2 | >2 | 0.5 | 0.125 | marolides, lincosamides, streptogramins (A) |
| AJUL10 | erythromycin | >2 | >2 | 1 | 0.125 | marolides, lincosamides, streptogramins (A) |
| AJUL11 | fusidic acid | 32 | >256 | 0.5 | 16 | fusidic acid |
| AJUL12 | mupirocin | >32 | >32 | 1 | 4 | mupirocin |
| AJUL14 | tetracycline | 16 | >64 | 2 | 8 | tetracyclines |
| AJUL15 | tetracycline | 8 | >64 | 2 | 8 | tetracyclines |
| AJUL16 | retapamulin | >0.5 | >0.5 | 0.125 | 0.0313 | pleuromutilins |
| AJUL18 | bacitracin | >512 | >512 | 32 | 16 | bacitracin |
| AJUL19 | ampicillin | 8 | 8 | 0.5 | 0.5 | β-lactams (penicillinase-susceptible) |
| AJUL20 | oxacillin | >8 | 4 | 1 | 0.5 | β-lactams (penicillinase-stable) |
| AJUL21 | fosfomycin | >512 | >512 | 128 | 32 | fosfomycin |
| AJUL22 | daptomycin | 16 | 4 | 4 | 1 | daptomycin |
| AJUL23 | rifampicin | >0.5 | >0.5 | 0.0078 | 0.0625 | rifamycins |
| AJUL24 | trimethoprim | >256 | >256 | 4 | >256 | diaminopyrimidines |
| AJUL25 | sulfamethoxzole | >2048 | >2048 | >2048 | >2048 | sulphonamides |
| AJUL26 | ciprofloxacin | 64 | 16 | 0.5 | 0.5 | fluoroquinolones |
| AJUL27 | novobiocin | >64 | >64 | 16 | 0.25 | aminocoumarins |
| AJUL28 | triclosan | 2 | 128 | 0.125 | 2 | triclosan |

**Table S2.** MIC values (% w/v) of Indigenous remedies' extracts tested against SH1000 cultured in SWF or MHB

| Extract code | MIC against SH1000 (% w/v) | |
|---|---|---|
| | SWF | MHB |
| Extracts that inhibited USA300 with higher potency in SWF than MHB | | |
| OEN1-022 | 0.415 | 0.415 |
| OEN1-067 | 0.225 | >0.225 |
| OEN1-053 | 0.341 | 0.341 |
| OEN1-063 | 0.231 | > 0.231 |
| OEN1-070 | > 0.015 | > 0.015 |
| Extracts that inhibited USA300 with relatively comparable potency in SWF and MHB | | |
| OEN1-058 | > 0.311 | 0.155 |
| OEN1-021 | > 0.269 | > 0.269 |
| OEN1-008 | 0.142 | 0.142 |
| OEN1-084 | 0.024 | 0.012 |
| OEN1-078 | 0.122 | 0.061 |
| OEN1-079 | 0.045 | 0.022 |
| OEN1-069 | 0.149 | 0.074 |
| OEN1-049 | 0.092 | 0.023 |
| OEN1-064 | 0.260 | 0.260 |
| OEN1-050 | > 0.150 | 0.075 |
| OEN1-011 | 0.344 | 0.344 |
| OEN1-036 | 0.141 | 0.141 |
